# The TonB dependent uptake of pyrroloquinoline-quinone (PQQ) and secretion of gluconate by *Escherichia coli* K-12

**DOI:** 10.1101/2022.06.07.495086

**Authors:** Klaus Hantke, Simon Friz

**Author notes:** Abbreviations: AUC, area under the curve in a mass spec; PQQ, pyrroloquinoline quinone; PTS, phosphotransferase sugar transport system.

## Abstract

Glucose is taken up by *Escherichia coli* through the phosphotransferase system (PTS) as the preferred carbon source. PTS mutants grow with glucose as a carbon source only in the presence of pyrroloquinoline quinone (PQQ), which is needed as a redox cofactor for the glucose dehydrogenase Gcd. The membrane-anchored Gcd enzyme oxidizes glucose to gluconolactone in the periplasm. For this reaction to occur, external supply of PQQ is required as *E. coli* is unable to produce PQQ *de novo*. Growth experiments show that PqqU (YncD) is the TonB-ExbBD dependent transporter for PQQ through the outer membrane. PQQ protected the cells from the PqqU dependent phage IsaakIselin (Bas10) by competition for the receptor protein. As a high affinity uptake system PqqU allows *E. coli* to activate Gcd even at surrounding PQQ concentrations of about 1 nmol/l. At about 30 fold higher PQQ concentrations the activation of Gcd gets PqqU independent. Due to its small size Pqq may also pass the outer membrane through porins. The PQQ dependent production of gluconate has been demonstrated in many plant growth promoting bacteria that solubilise phosphate minerals in the soil by secreting this acid. Under Pi limiting conditions also *E. coli* induces the glucose dehydrogenase and secretes gluconate, even in absence of PTS, that is, even when the bacterium is unable to grow on glucose without PQQ.

## Introduction

Enterobacteria like *Escherichia coli* use glucose as the preferred carbon source. The uptake depends on the phosphotransferase (PTS) system, which is required for translocation of the carbohydrate across the inner membrane concomitant with its phosphorylation. However, glucose may also be oxidized in the periplasm to gluconolactone by the glucose dehydrogenase Gcd (van Schie et al., 1985). In addition, the periplasmic soluble aldose sugar dehydrogenase Asd (YliI), which has a very broad substrate specificity, may contribute to glucose oxidation, leading to formation of gluconate (Southall et al., 2006). Gluconate may then be transported into the cell by one of the four sugar acid transporters GntU, GntT, GntP, or IdnT(=GntW) (Peekhaus et al., 1997) for further catabolism. However, gluconate that is produced in the periplasm may not always be taken up directly into the cytoplasm. It is also found in the medium as reported by (Bouvet and Grimont, 1988). Both dehydrogenases Gcd and Asd require pyrroloquinoline-quinone (PQQ) (Fig.1) as redox cofactor for their activity. Since *E. coli* is unable to produce PQQ (Matsushita et al., 1997), this sugar oxidation is not observed without exogenous addition of PQQ. However, in the context of a microbiome where PQQ may be synthesized by other Gram negative bacteria like *Klebsiella*, *Pseudomonas*, *Acinetobacter*, this kind of sugar utilisation may be important for *E. coli*. The secretion of large (µmol/l) amounts of PQQ into the growth medium has been demonstrated for more than 15 different soil bacteria (McIntire, 1998).

**Fig.1:**
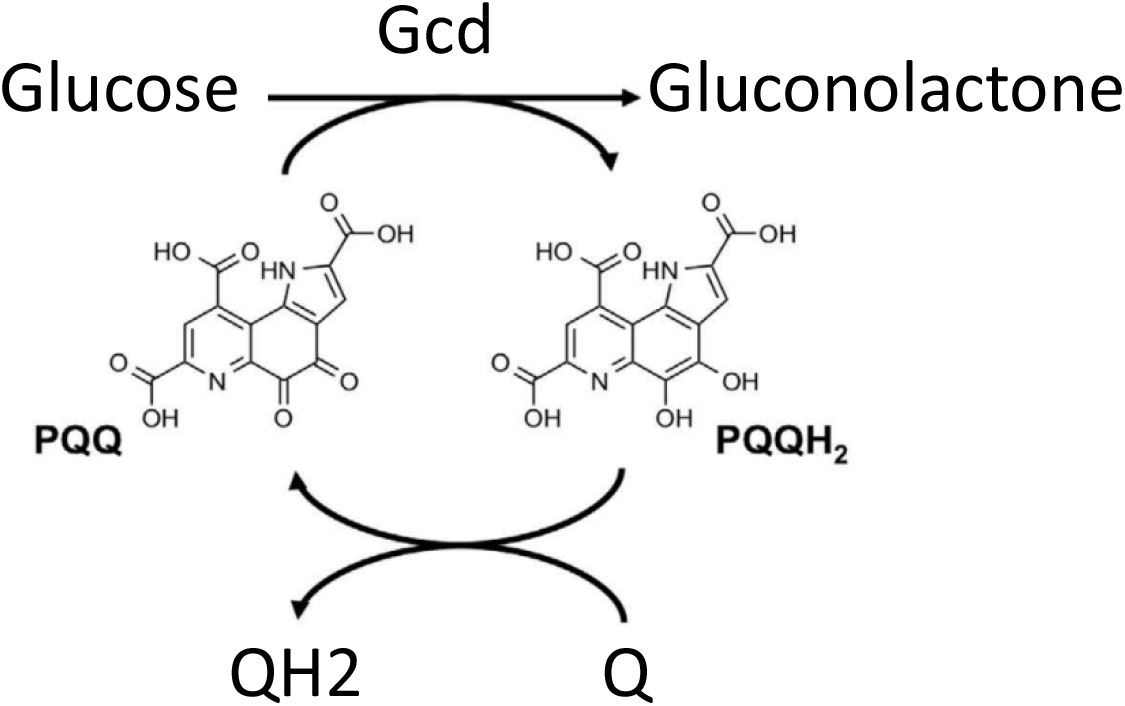
Overview of the PQQ/Gcd-dependent oxidation of glucose in gluconolactone. Pyrroloquinoline quinone (PQQ, 4,5-dihydro-4,5-dioxo-1*H*-pyrrolo[2,3-*f*]quinolone-2,7,9-tricarboxilic acid, methoxatin) is a redox cofactor for quinoproteins like glucosedehydrogenase (Gcd), PQQH_2_ may be oxidized by ubiquinone (Q) (Cordell and Daley, 2022).

The complicated biosynthesis of PQQ has been elucidated in the recent years. In a ribosomally synthesized peptide, a glutamate residue is linked to a tyrosine residue and further shaped to PQQ (Zhu and Klinman, 2020). It may be expected that PQQ is present in very low amounts in the natural environment of *E. coli.* Rare substances like vitamin B12 and iron siderophores are generally imported across the *E. coli* outer membrane by TonB dependent receptors. These uptake processes are energized by the membrane potential via the TonB-ExbBD complex which allows high affinity uptake (Braun, 1995). Of the predicted nine TonB-dependent receptors in *E. coli* K-12, six are known to mediate the uptake of iron siderophores (enterochelin and its degradation products, ferrichrome, coprogen and iron-citrate) (Braun and Hantke, 2007). The TonB dependent protein BtuB is the receptor for vitamin B12 (Bradbeer, 1993), which is used as a cofactor for methylgroup transfer reactions and methionine biosynthesis. The structure of the two other predicted TonB dependent proteins YncD and YddB has been elucidated, but their function in *E. coli* is still unknown (Grinter and Lithgow, 2020). Upstream of *yncD* on the complementary strand the gene *yncE* encodes a periplasmic protein that has been shown to bind DNA and whose structure has been determined to be a seven bladed propeller (Kagawa et al., 2011). In quinoproteins PQQ is often bound to bladed propeller structures. Therefore, this protein is predicted in some databases as a PQQ binding protein (Keseler et al., 2021), although its function is unknown. Here, we tested whether PQQ is a substrate of YncD. Since we found evidence that YncD mediates uptake of PQQ, we propose to name this receptor PqqU based on its function.

## Results

### YncD is the receptor for PQQ

Glucose is mainly taken up by the PTS system for further metabolism. In a PTS deficient mutant glucose is oxidized in the periplasm to gluconolactone by the glucose dehydrogenase Gcd (Cleton-Jansen et al., 1990) and the sugar aldose dehydrogenase YliI (Southall et al., 2006). Gluconolactone may hydrolyse spontaneously or, more likely, is hydrolysed to gluconate by an up to now unknown lactonase. The product of the hydrolysis, gluconate, is then taken up and further metabolised. PQQ dependent growth of *E. coli* 6281 Δ(*ptsHI crr*) (abbreviated Δ*pts*) was demonstrated in liquid minimal medium MOPS or M63, containing 0,1 mmol/l or 100 mmol/l Pi, respectively, supplemented with 0,1% glucose and different concentrations of PQQ (Fig.2). Even at a concentration of 1 nmol/l PQQ stimulated rapid growth of 6281 Δ*pts* in both minimal media (Fig.2A,C). In contrast, at low PQQ concentrations the *E. coli pts pqqU* deletion mutant displayed a delayed growth start in the MOPS medium, an effect that was concentration dependent and not observed at high PQQ concentrations (Fig.2B). In the M63 medium, the Δ*(ptsHI crr)* Δ*pqqU* deletion mutant showed a long lag phase, and at lower PQQ concentrations no growth was observed in the time given (2D). When instead of glucose, gluconate was used as carbon source, both the PTS deficient strain and the PTS and PqqU deficient strain were able to grow without addition of PQQ (Fig.2). This shows that only the oxidation of glucose to gluconate is affected in the absence of PqqU activity.

**Fig.2:**
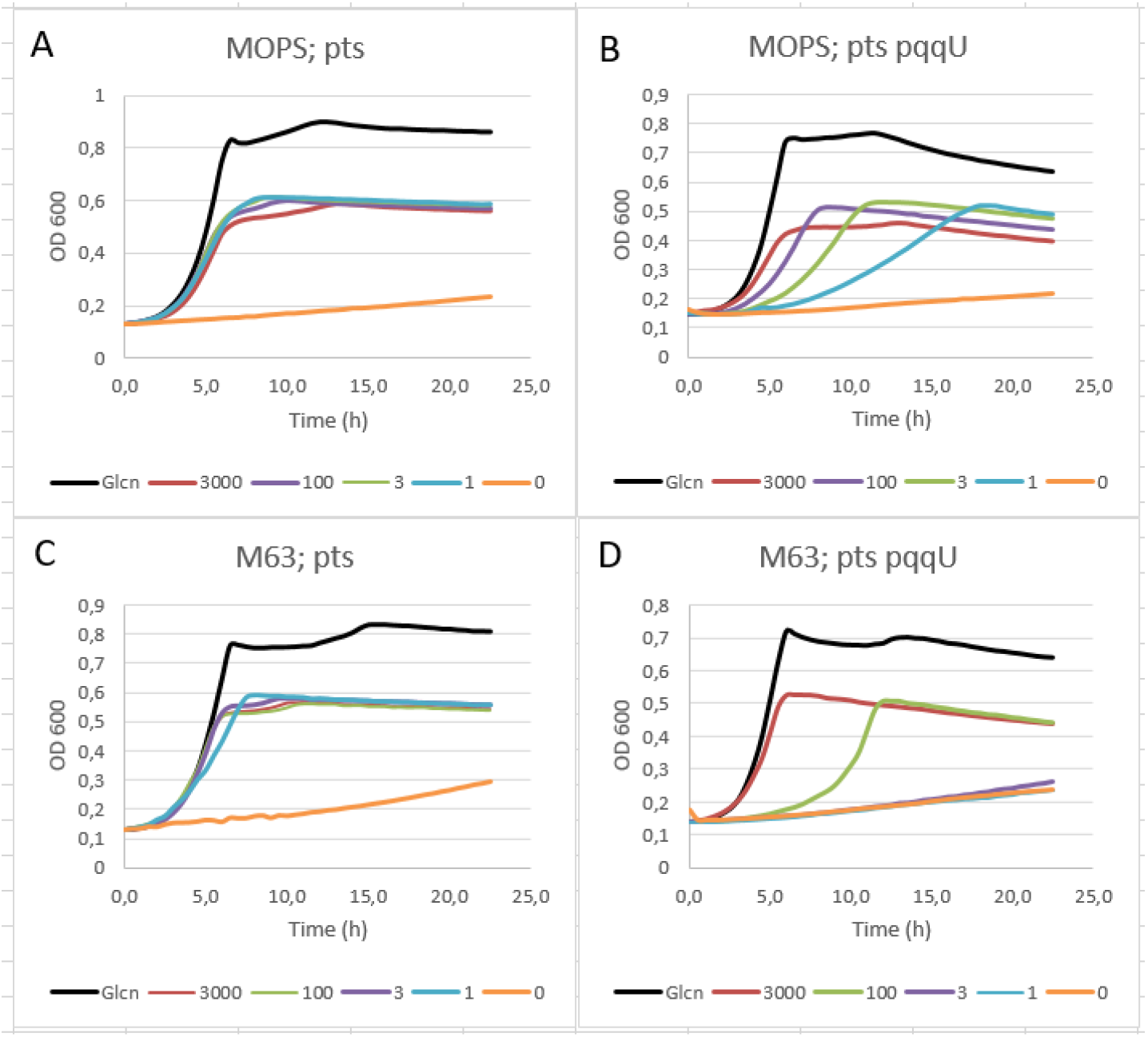
PQQ dependent growth of a PTS deficient *E. coli* strain. Cells were pregrown in minimal medium M63 0,6% glycerol over night. Strains 6281 Δ*(ptsHI crr*) (abbreviated *pts*) (A, C) and 6289 Δ*(ptsHI crr*) Δ*pqqU* (abbreviated *pts pqqU*) (B, D) were grown in minimal medium MOPS 0,1 mmol/l Pi or M63 100 mmol/l Pi as indicated in the chart title in a 96 well plate. Carbon source was 0,2 % gluconate (Glcn, black) or 0,1 % glucose. The different concentrations of PQQ, 3000 to 0 nmol/l, are indicated in the bottom line of the graph.

Hydrophilic molecules with a molecular weight below 600 Da are expected to pass through the outer membrane by diffusion across porins (Nikaido, 2003). Growth of the YncD deficient mutant at high PQQ concentrations may be explained by the small size of PQQ which has a molecular weight of 330 Da and can freely diffuse across porins, and thus fulfil the demands of the glucose dehydrogenase Gcd as a cofactor. At low concentrations the high affinity uptake system for PQQ is necessary.

As a TonB-dependent transport is predicted from the structure of PqqU, we tested the growth of a PTS and TonB deficient mutant, *E. coli* 6284 Δ(*ptsHI crr*) *tonB*. In the presence of PQQ a growth response to 3000 nMol/l PQQ was observed, but no response to 100nmol/l PQQ (Fig.3A). Since the energisation of these receptors is dependent on the TonB/ExbB/ExbD complex an *exbB*::Tn10 mutant was also tested. In minimal medium with gluconate, the ExbB mutant grew as well as the parent strain. Growth with glucose was stimulated by high PQQ concentrations but delayed at lower ones (Fig.3B) indicating a dependence of PQQ uptake on ExbB. In summary, at low concentrations *E. coli* appears to require the concerted activity of PqqU and TonB/ExbBD for PQQ uptake.

**Fig.3:**
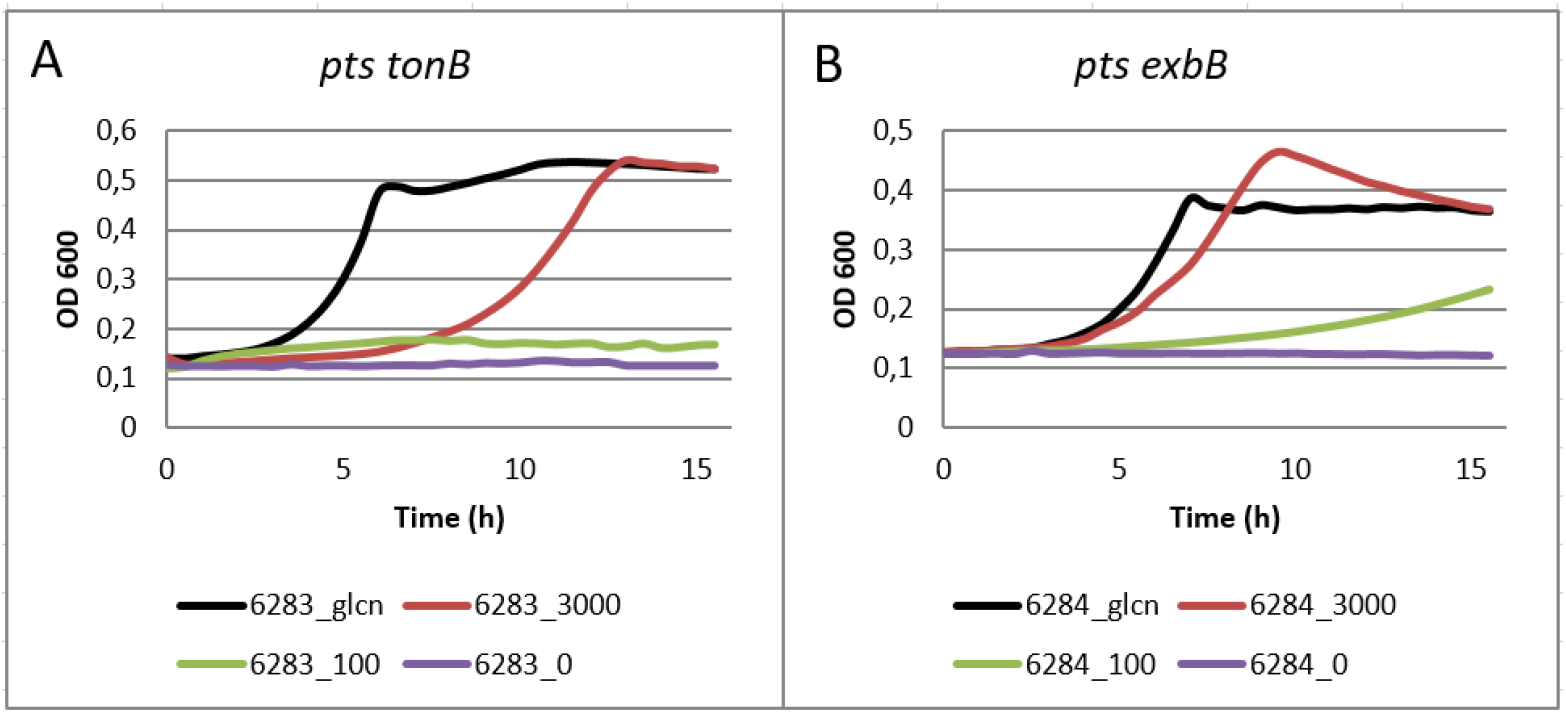
Growth of the *E. coli pts* deletion mutant in the presence of low concentrations of PQQ depends on the TonB/ExbBD system. Growth of TonB and ExbB mutants on M63 minimal medium with 0,1% gluconate (glcn) or 0,1% glucose supplemented with 3000, 300, 30, 10 or 0 nmol/l PQQ. Strains: (A) 6284 Δ(*ptsHI crr*) *tonB*, (B) 6283 Δ(*ptsHI crr*) *exbB*

### Inhibition of phage infection by PQQ

The phages T5 and BF23 are known to use the TonB dependent outer membrane proteins FhuA and BtuB, respectively, as receptors for infection. It was shown that the substrates of these receptors, ferrichrome and vitamin B12, respectively, could protect the cells from infection (Bradbeer et al., 1976, Hantke and Braun, 1978). Since there are also phages known to use PqqU as a receptor, such as phage IsaakIselin (Bas10) (Maffei et al., 2021), we asked whether there could be competition between the substrate PQQ and phage for binding to PqqU. Plating efficiency of Bas10 was the same on the parental strain BW15113 and the TonB deficient mutant JW5195 while the PqqU mutants JW1446 and 6289 were resistant. On agar plates the application of filter paper discs soaked with PQQ inhibited plaque formation by Bas10 (Fig.4). Interestingly, the single *tonB* and the double *tonB pts* mutants were protected more efficiently than the parental strains as a roughly 1000-fold higher concentration of PQQ was necessary to protect the latter. These findings are in line with what has been observed for the ferrichrome and vitamin B12 uptake systems: due to the lack of TonB PQQ remains bound to the receptor and efficiently blocks phage adsorption.

**Fig.4:**
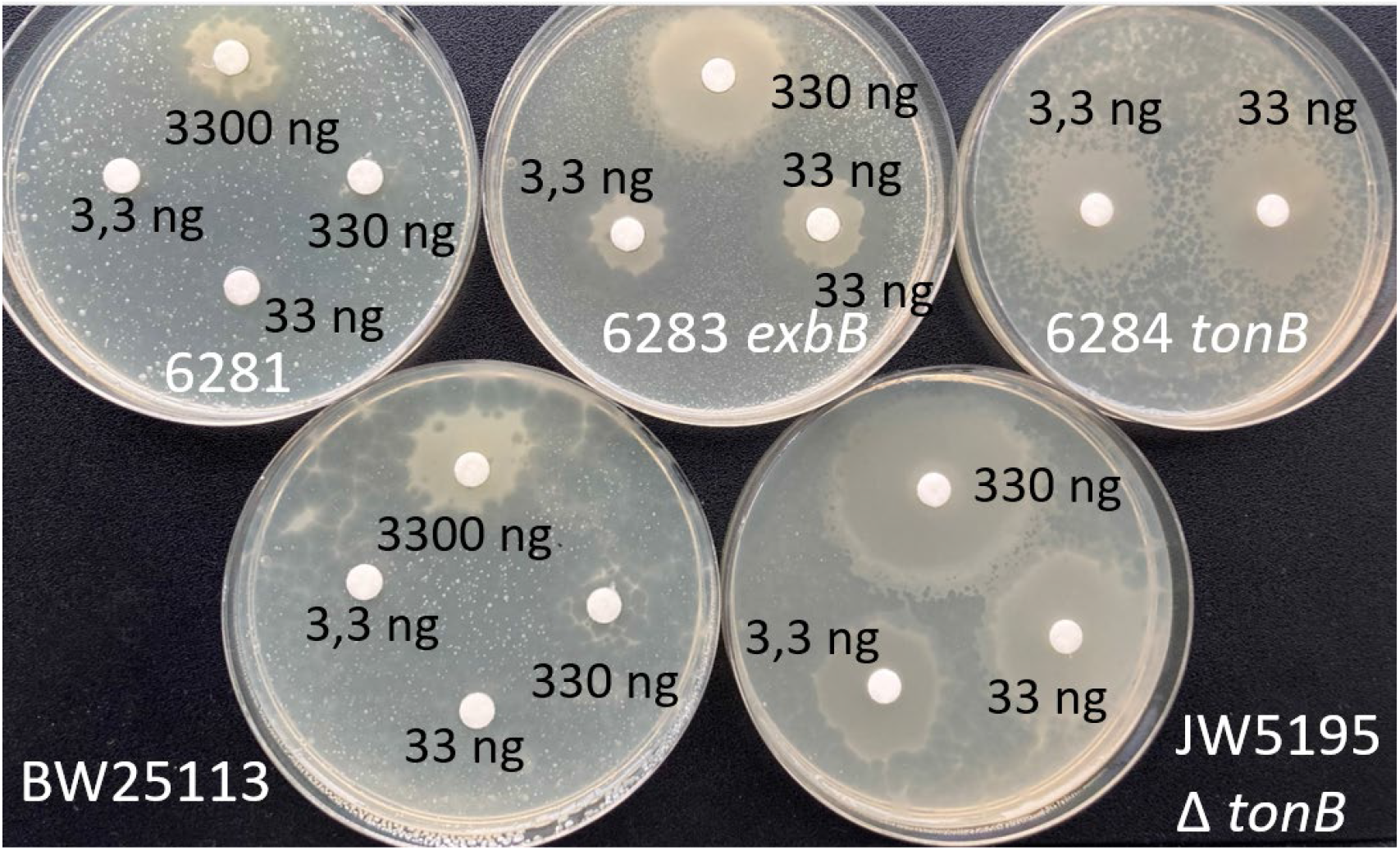
Inhibition of phage infection by PQQ. An LB agar plate was overlaid with 3ml LB soft agar containing about 5×10^8^ cells of the indicated bacterial strains and 10^4^ Bas10 phages. Filter paper discs with the indicated amount of PQQ were applied. After incubation at 37°C bacteria protected from the phage induced lysis were able to grow in the PQQ diffusion zone around the paper discs. In the TonB mutants the substrate is not taken up, remains bound to the receptor and inhibits adsorption of the PqqU dependent phage Bas10. The ExbB mutant is less well protected due to the partial complementation by *tolQR*.

### Influence of phosphate on PQQ dependent growth and secretion of the oxidation product

As it can be seen in Fig.2 there is a dramatic difference in the growth response of the PqqU mutant between a phosphate rich and phosphate poor medium. Adamowicz et al. (1991) studied two glucose negative mutants of *E. coli*, *ptsM ptsG* and *pgi zwf* respectively, which were unable to grow on glucose in a minimal medium. Only after addition of PQQ growth was observed. Both strains had a rather long lag phase when they were pre-cultured in a phosphate rich medium. It was suggested that the porin PhoE, with its known preference for anionic substrates, could be the gate for PQQ to get access to the inner membrane-anchored glucose dehydrogenase Gcd facing the periplasm (Fig.8). Pre-growth of *E. coli* in a phosphate rich medium would repress PhoE expression and lack of PhoE would slow down PQQ uptake. However, this hypothesis was not experimentally verified.

Phosphate limitation leads to a general nutrient limitation with major changes in gene expression, which are mediated by the regulatory protein DksA (Gray, 2020). Using an RNA sequencing screen, Gray (2020) analysed those changes after a downshift from LB medium to a MOPS 0.1 mmol/l Pi medium. Interestingly, *gcd* expression increased about 10-fold (Gray, 2020). This could lead to a different interpretation of the data of Adamowicz *et al*. (1991). Cells grown in a phosphate rich medium may have Pi reserves and may need more time to adapt to the stress imposed by low phosphate levels. In addition, we did not observe an influence of PhoE on PQQ dependent growth (data not shown).

Among plant growth promoting bacteria like *Pseudomonas*, PQQ-dependent glucose oxidation and secretion of gluconate is widespread (Miller et al., 2010, Raymond et al., 2021). Gluconate solubilises phosphate minerals and in doing so furthers plant growth. Not only *Pseudomonas* but also Enterobacteria such as *Serratia* (Luduena et al., 2017) and *Klebsiella* (Hommes et al., 1989) strains are described to produce PQQ and secrete gluconate.

We investigated whether *E. coli* K-12 also secretes the oxidation product of Gdh into the medium. Strain BW25113 was grown with PQQ in MOPS (0,1mmol/l Pi) medium with 0,1 % glucose. The supernatant was analysed by liquid chromatography-mass spectrometry (LC-MS). About 3 mmol/l gluconate were found during the late logarithmic growth phase in the medium (Fig.5). This was consumed in the late logarithmic growth phase. Instead, much less gluconate (less than 0,2 mmol/l) was secreted in the phosphate rich M63 medium. Interestingly, also the *pts* deletion mutant H2681 secreted gluconate into the medium, although one would have expected gluconate to be directly transported from the periplasm to the cytoplasm. In MOPS medium with 0,1% glucose, about 0,9 mmol/l gluconate was found in the late log phase. In M63 a similar amount of about 0,7 mmol/l gluconate was determined. It remains to be clarified how much gluconolactone/gluconate is directly taken up from the periplasm.

**Fig.5:**
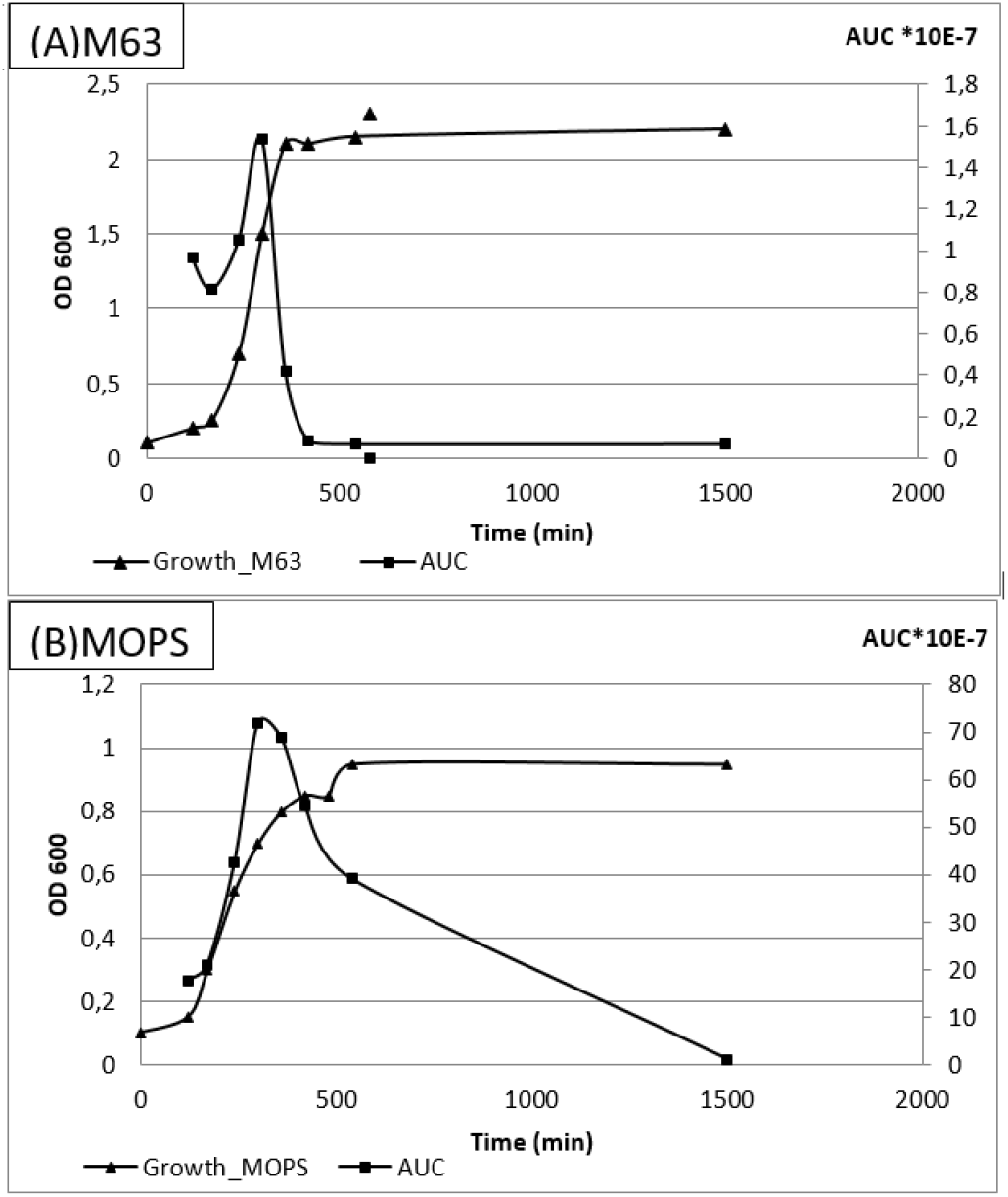
Gluconate production of BW25113. *E. coli* K-12 BW25113 was grown in minimal medium M63 (A) or MOPS (B) 0,1 mmol/l Pi with 0,1 % glucose and 0,6 µmol/l PQQ. Gluconate in the supernatant was determined by LC-MS; 70*10^7^ AUC (area under the curve in a mass spec) are about 3 mmol/l gluconate; 1,5*10^7^ AUC are below 0,2 mmol/l gluconate. Note the different scales of the abscissa (AUC). Cells grown in M63 medium achieved twice the cell density compared to cells grown in MOPS.

### Visualisation of gluconate secretion

Secretion of acids may be visualised on Mac Conkey agar plates. On glucose-containing solid medium *E. coli* forms red colonies because glucose overflow metabolism causes secretion of organic acids such as acetic acid. The acidification leads to a colour change of the pH indicator neutral red from yellow to red, and to the precipitation of bile acids which makes the agar turbid. PTS mutants cannot utilise glucose and form pale whitish colonies. Application of filter paper discs soaked with PQQ allowed *E. coli* Δ*pts* to utilise glucose. This resulted in red colonies and a red turbid halo in the diffusion zone. The size of the red and turbid zones was dependent on the PQQ concentration (Fig.5A). However, inoculum size and incubation time had a large influence on the size of the halo. While for the *E. coli* Δ*pts* application of about 10 nmol/l PQQ in the agar led to the formation of red colonies with a turbid halo, for strain 6289 Δ*pts* Δ*pqqU*, an approximately 30 to 100 fold higher PQQ concentration was necessary to observe red colonies with a turbid halo (Fig.5A). This again indicates that the *pqqU* mutant lacks a high affinity uptake system for PQQ.

A similar difference in colony colour development was observed when Mac Conkey plates contained defined PQQ concentrations (Fig.5B). Colonies of 6289 *pqqU* null mutant were only red with 1000 nmol/l PQQ while the parent strain 6281 produced even at 3 nmol/l PQQ red colonies. In addition, an YncE deficient mutant was tested, which in both this plate assay and in growth experiments (data not shown) behaved like the parent strain.

The addition of 10 µmol/l iron to the Mac Conkey agar plates was necessary to suppress the acid production of strain 6284 Δ*pts* Δ*tonB* with and without glucose (Fig.5A and Fig.S1). This indicates that acid is produced from peptones. After addition of iron, pale white colonies appeared (Fig.S1). The glucose-independent acid production is a consequence of the *tonB* mutation. The strain is unable to utilise siderophore bound iron and secretes the high affinity iron chelator enterochelin to overcome the iron deficit. This aggravates the situation as more iron is chelated and less iron is available for low affinity uptake systems. It seems that acids are secreted due to the internal iron deficit. In chemostat cultures upon iron limitation an overflow metabolism with secretion of acetate and lactate has been observed in *E. coli* (Folsom et al., 2014). Interestingly, a similar observation has been made in *Pseudomonas* (Sasnow et al., 2016).

On iron supplemented Mac Conkey plates with glucose (Fig.5A), the TonB mutant secretes acids only at relatively high PQQ concentrations demonstrating that PQQ is not taken up by the high affinity uptake system.

The ExbB deficient mutant showed even larger red turbid zones around the PQQ discs in comparison to the parent strain. This may be explained by the partial complementation of ExbBD by TolQR which is observed in several TonB dependent uptake systems (Bradbeer, 1993, Braun and Herrmann, 1993). The lower velocity of PQQ uptake may allow further diffusion of PQQ, especially at low cell densities and could explain the larger response zone.

### PQQ has no direct influence on the *E. coli* iron metabolism

The crystal structure of PqqU (YncD) has been solved (Grinter and Lithgow, 2020) and a certain similarity to the di-iron-di-citrate siderophore receptor FecA was observed. The predicted binding site contains several positively charged residues. This argues for a negatively charged substrate, but an iron-citrate complex did not fit. PQQ (Fig.1) fulfils the postulate of the authors “…the substrate of YncD is likely to be relatively small and negatively charged.” (Grinter and Lithgow, 2020). PQQ docking experiments by Vikram Alba showed that the pocket is large enough to accommodate PQQ. However, the docking method used identified 9 different poses (Fig.7).

**Fig.6:**
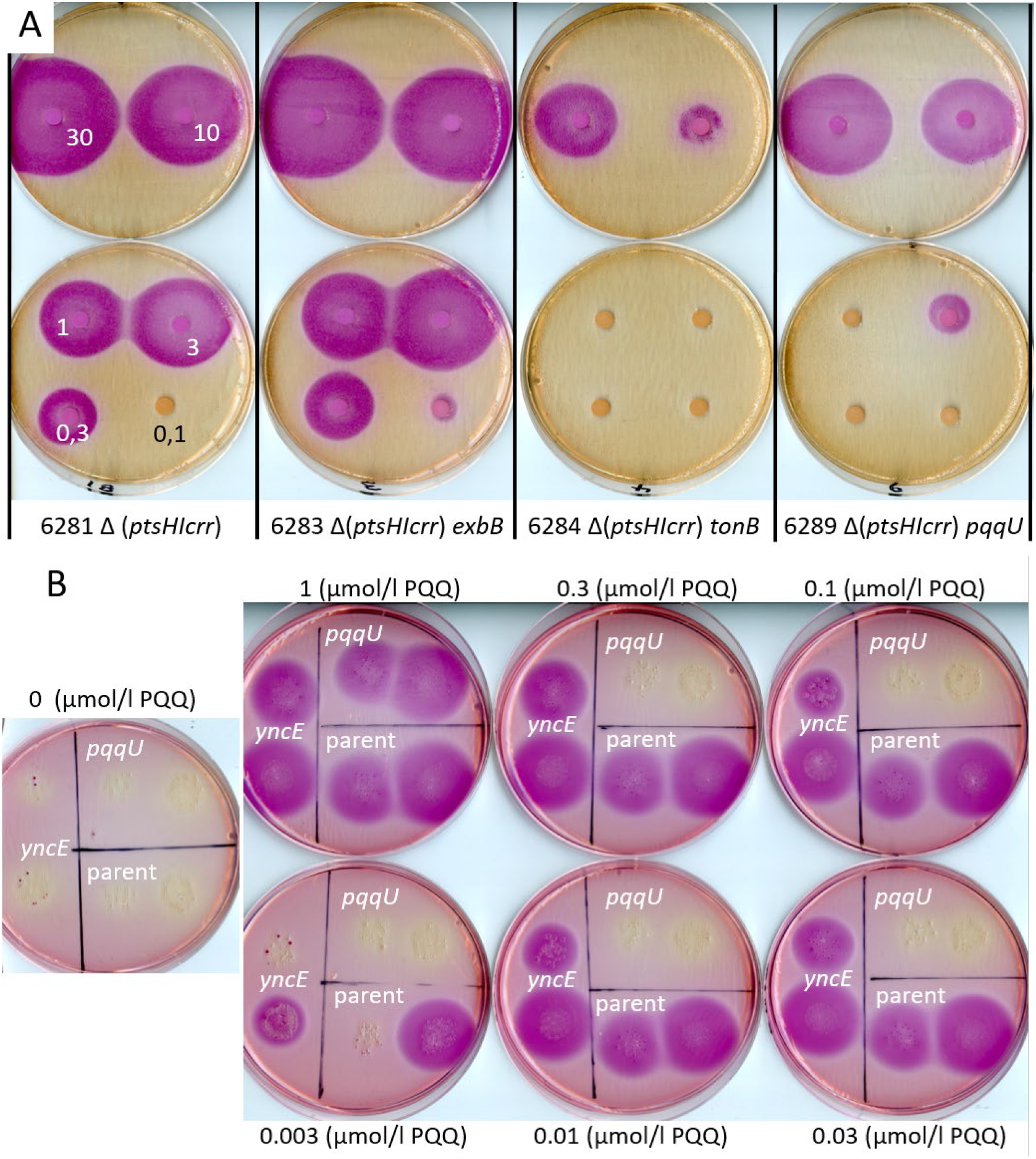
The PqqU/TonB/ExbB uptake system is required for PQQ-dependent production and secretion of gluconate at low PQQ concentrations. (A) Approximately 10^4^ cells of the indicated strains were spread on MacConkey agar plates (0,1 % glucose 10 µmol/l Fe(III)Na(EDTA)) and filter paper discs containing 30 ng to 0,1 ng PQQ were applied. Plates were incubated at 37 °C for about 14 hours. (B) MacConkey glucose agar plates were prepared containing 0 to 1µmol/l PQQ as indicated. 20µl containing about 30 to 300 colony forming units of the strains 6281 *Δpts,* 6289 *Δpts ΔpqqU* and 6280 *Δpts ΔyncE* were applied. After about 18h of incubation at 37°C, pictures were taken.

**Fig.7:**
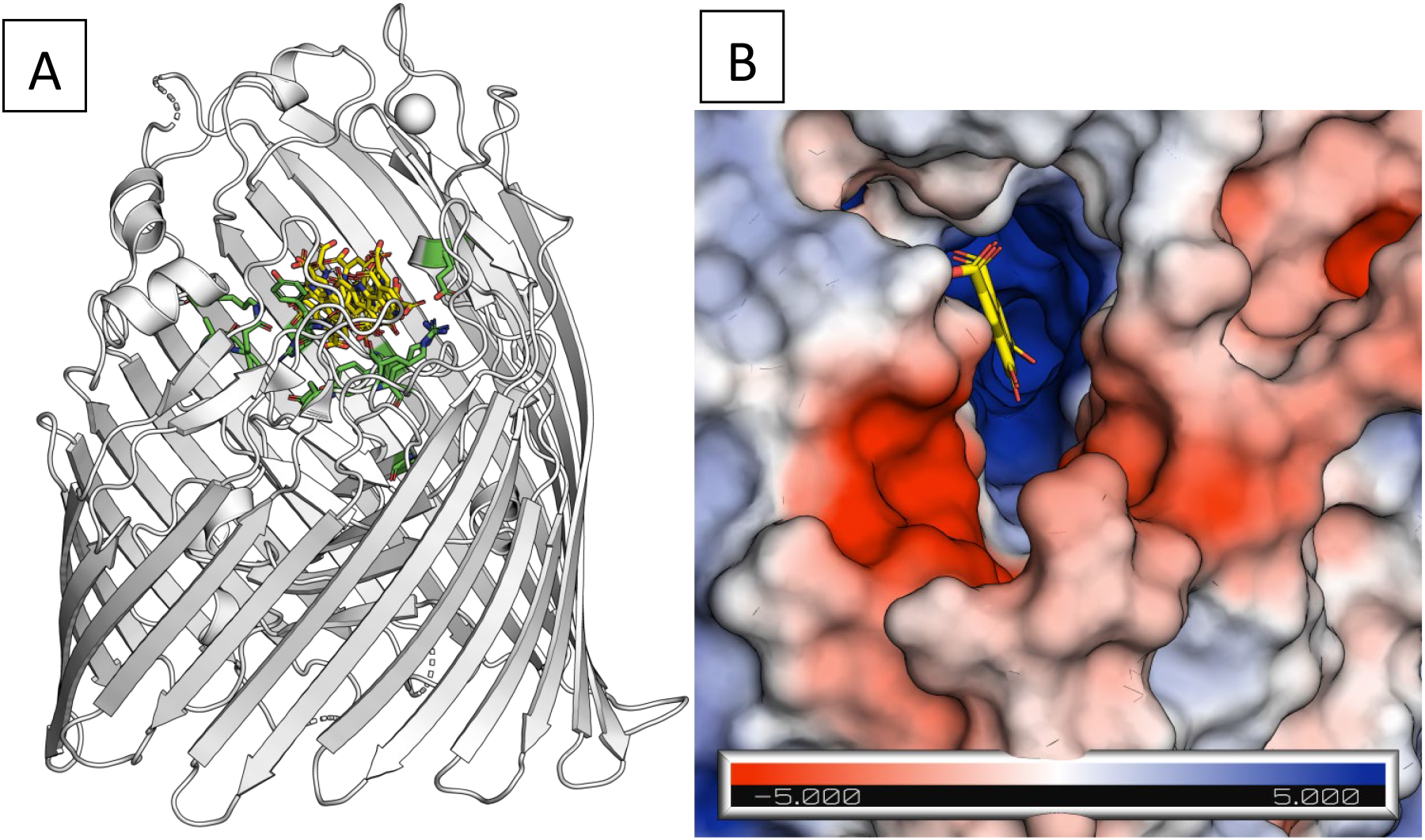
Crystal structure of PqqU (YncD) (Grinter and Lithgow, 2020) (A) with 9 different poses of PQQ in the binding pocket. (B) The best scoring pose in the binding pocket is shown.

The carboxylate groups of PQQ are known to interact with metals like Ca2+, Mg2+ and lanthanoids (Wehrmann et al., 2020) in the binding pockets of quinoproteins. Therefore, we tested whether the supply of PQQ in an iron poor environment would influence the iron balance in the cells. In an iron rich medium the *fhuF-lacZ* fusion of strain H1717 is repressed by the iron uptake regulator Fur (Stojiljkovic et al., 1994). Supply of PQQ did not change the expression of the *fhuF-lacZ* fusion neither under iron rich nor under iron poor conditions (data not shown). This indicated that PQQ neither increased nor decreased the internal iron level. These results are in line with the findings of different groups that PqqU is not regulated by Fur and does not transport iron in relevant amounts (Grinter and Lithgow, 2020).

## Discussion

Vitamin B12 and PQQ are needed as cofactors for certain enzymes. Since *E. coli* cannot synthetize them *de novo*, the bacterium depends on an external supply. In the environment, both substances may be present only in low concentrations. Accordingly, uptake of vitamin B12 requires the high affinity, TonB dependent receptor BtuB (Bradbeer, 1993). It is not surprising that also PQQ is taken up by a TonB dependent receptor, namely PqqU. Unlike vitamin B12, PQQ is a small molecule and may also pass the outer membrane without PqqU. However, this diffusion dependent uptake process is rather inefficient, which may be a disadvantage when only low amounts of PQQ are present.

As a “common good”, PQQ like vitamin B12 may promote interspecies interactions in the microbiome (Sokolovskaya et al., 2020). PQQ dependent gluconate production is characteristic for numerous plant growth promoting bacteria (Shen et al., 2012, Meyer et al., 2011, Wu et al., 2022). Organic acids like gluconate mobilise phosphate from insoluble calcium phosphates and thereby further plant growth. There is evidence that phosphate limitation induces PQQ synthesis in *Pseudomonas putida* (An and Moe, 2016) and *Serratia marcescens* (Luduena et al., 2017). Interestingly, iron limitation also led to a high secretion of gluconate (44% of the supplied glucose) in *Pseudomonas putida* (Sasnow et al., 2016). However, under the growth conditions used, the additionally synthesized siderophore mobilised most of the iron mineral supplied as goethite. Gluconate did not contribute significantly to the iron supply of the cells (Sasnow et al., 2016).

In *E. coli*, gluconate secretion has been described for a *pgl* mutant, which is unable to produce phospho-gluconolactonase in the cytoplasm (Kupor and Fraenkel, 1969). For the product of Gcd, gluconolactone, it is unknown whether it is enzymatically hydrolised. Under physiological conditions gluconolactone is rather unstable and may hydrolise spontaneously before or after it is secreted. A gluconolactonase in the periplasm was identified in a gluconate secreting *Pseudomonas* (Tarighi et al., 2008). Since gluconolactones are toxic due to their acylating activity, a lactonase may be expected for “house cleaning” purposes (Galperin et al., 2006). A possible candidate could be YncE, which is encoded upstream of *pqqU* on the complementary strand. The bladed propeller structure of YncE (Kagawa et al., 2011) is often found in quinoproteins. Therefore, in the beginning of our studies we had the hypothesis that it could be a PQQ binding protein involved in transport. However, PQQ-dependent growth of the *yncE* deletion strain did not differ from that of the parent strain (Fig.5B and data not shown). This would not necessarily be expected for a gluconolatonase mutant, since a *pgl* mutant shows only minor growth defects with glucose as carbon source (Kupor and Fraenkel, 1972). Sequence comparison did not allow assignment of a function for *yncE*. In this context, it is interesting to mention that YncE has been found to be a protective antigen in *Escherichia* (Moriel et al., 2016) and a diagnostically interesting antigen in *Salmonella* (Franklin et al., 2020). Also PqqU (= YncD) from *Salmonella* has been demonstrated in mice to be a protective antigen (Xiong et al., 2012).

The importance of PQQ for *E. coli* has also been demonstrated by chemotaxis towards PQQ (de Jonge et al., 1996). However, it may not be the molecule PQQ that is detected in this case, but the redox state of the respiratory chain (Samanta et al., 2016) which may determine the direction during swimming.

## Conclusion

PQQ may be taken up with the help of the TonB-ExbBD energized outer membrane receptor PqqU (YncD) (Fig.8). In the presence of PQQ the quinoprotein Gcd oxidises glucose to gluconolactone. Where and how gluconolactone is hydrolised, enzymatically or spontaneously, is unknown. Appreciable amounts of gluconate appear in the medium and are completely metabolised in a minimal medium when the stationary growth phase is reached.

**Fig.8:**
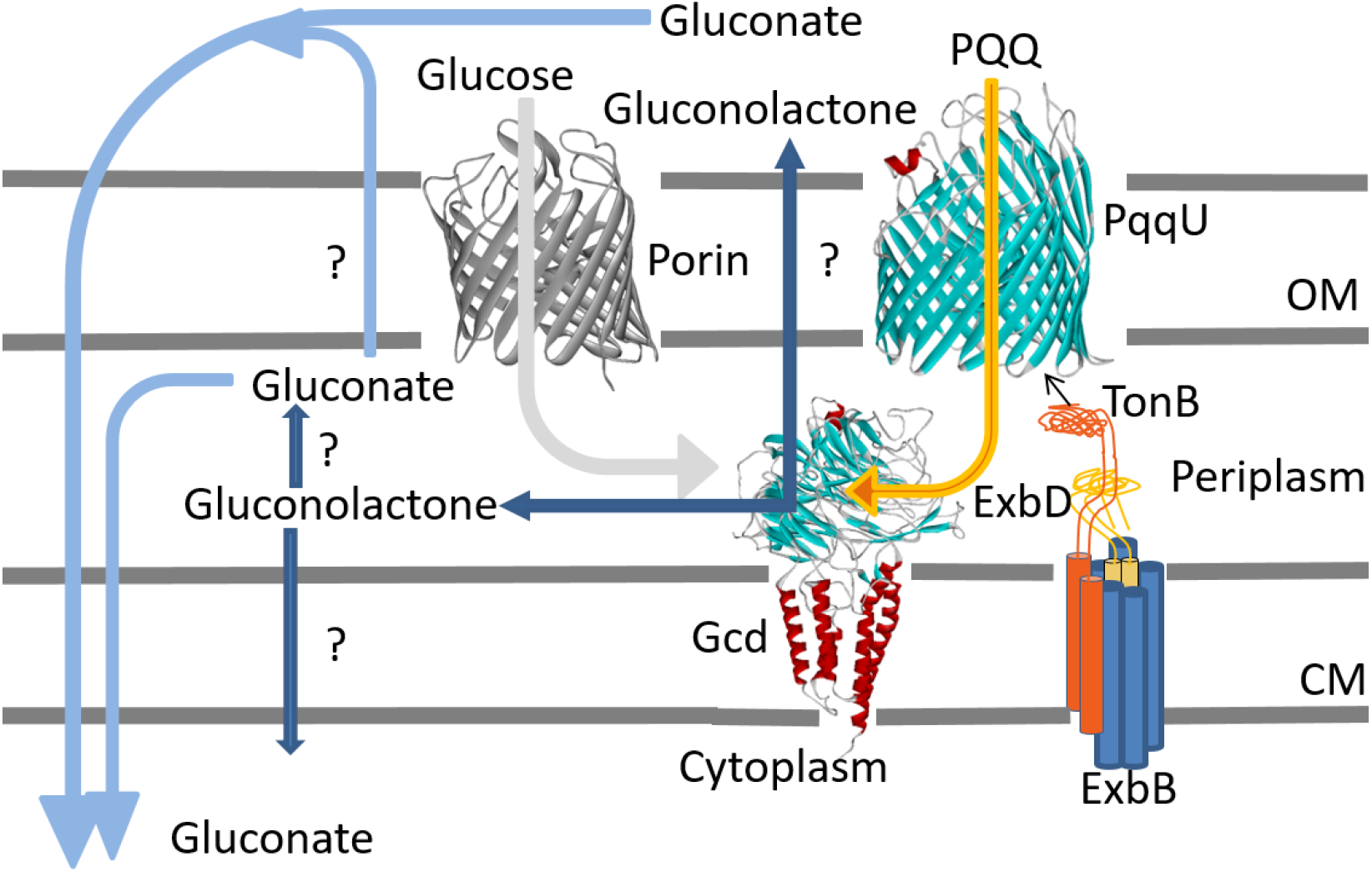
PQQ uptake and gluconate secretion in *E. coli*: The question marks indicate that it is unknown where and how gluconolactone hydrolises or is enzymatically hydrolised.

## Materials and methods

### Bacterial growth conditions and strains

Bacteria were grown in LB or Difco Nutrient Broth at 37°C, with kanamycin (50µg/ml) or tetracycline (15µg/ml) if necessary. Difco Mac Conkey agar base with 10g/l glucose was used and supplemented where indicated with 10 µmol/l Fe(III)Na-EDTA. Approximately 10 000 bacteria per plate were applied to demonstrate PQQ-dependent sugar utilisation. The diameter of the red zone was determined after 18 h.

The minimal medium M63 (containing per liter: 5,3 g KH_2_PO_4_, 13,9 g, K_2_HPO_4_x3H_2_O, 2 g (NH_4_)_2_SO_4_, 1mM MgSO_4_ was used for growth. Carbon sources were added as indicated. OD_600nm_was measured in a TECAN SPARK 10M reader. The MOPS minimal medium contained 40 mM MOPS, 86 mM NaCl, 9,5 mM NH_4_Cl, 13,3 mM KCl; pH was adjusted to 7,2 with KOH. The Pi-limiting MOPS minimal medium contained 0,1 mM Pi by adding 1 ml of M63 salts/l. Carbon sources were added as indicated.

The bacterial strains used are listed in Table 1. Strains of the Keio collection (Baba et al., 2006) were kindly supplied by Heike Brötz-Oesterhelt. The kanamycin resistance gene was removed with pCP20 as described (Datsenko and Wanner, 2000) and further strain constructions were done by P1 transductions.

**Table 1:**
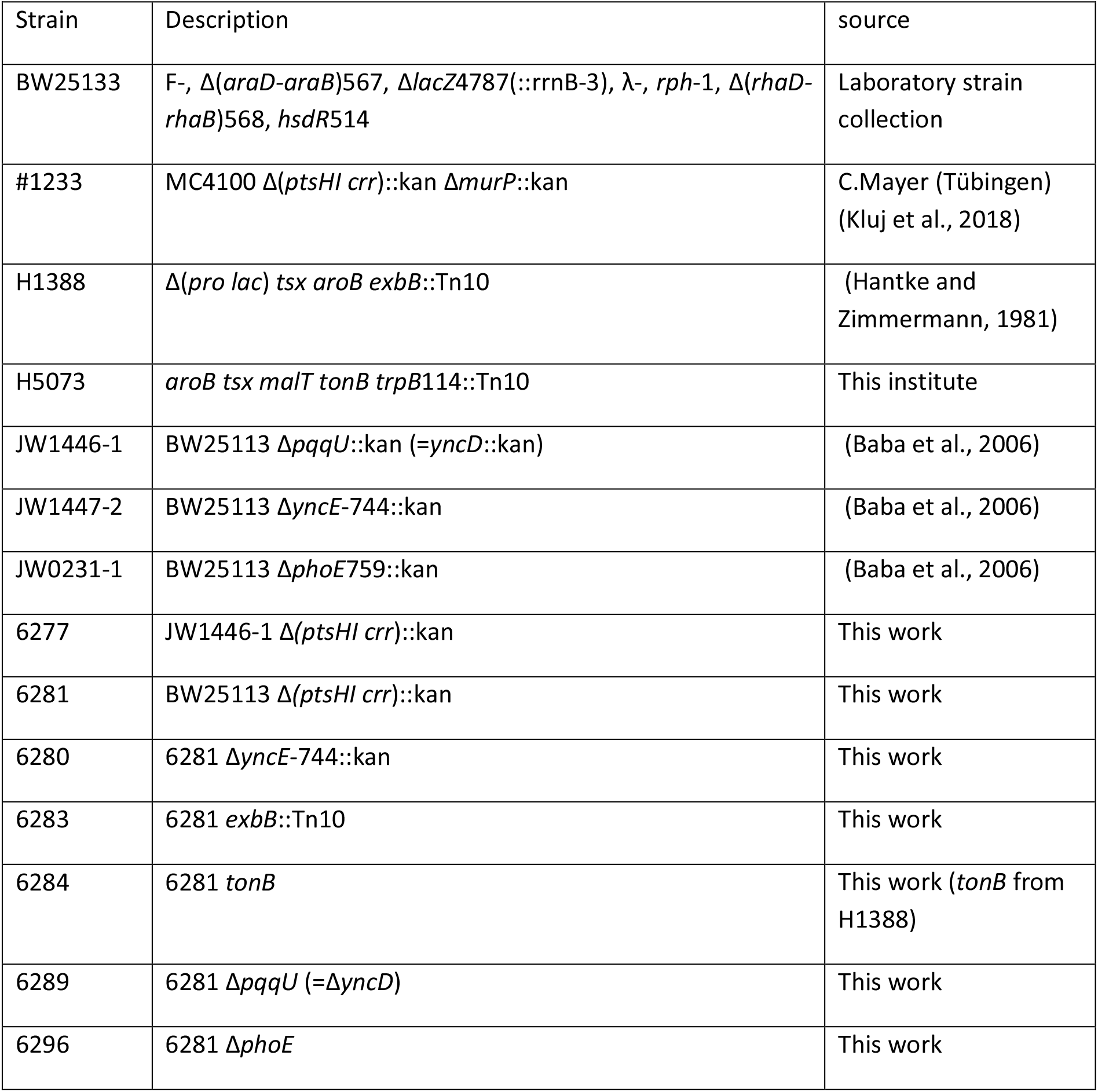
List of *E.coli* strains

Phage IsaakIselin (Bas10) was obtained from A. Harms (Basel) (Maffei *et al*.2021) and propagated on LB agar plates with BW25113 as host.

### LC-MS analysis

Bacteria were grown in baffled 100 ml Erlenmeyer flasks with 15 ml medium under constant shaking at 37°C. Samples were prepared with Costar® Spin-X® centrifuge filters immediately after drawing 0,5 ml from the culture supernatant. Extracellular gluconate was analysed using an electrospray ionization time of flight (ESI-TOF) mass spectrometer (MicrOTOF II; Bruker Daltonics), connected to an UltiMate 3000 high-performance liquid chromatography (HPLC) system (Dionex). Separation in the HPLC was carried out using a SeQuant ZIC-pHILIC column (PEEK 150 × 2,1 mm, 5 μm, 110 Å, Merck) at 30 °C with an CH_3_CN (buffer A) and 100 mM (NH_4_)_2_CO_3_, pH 9 (buffer B) buffer system. 5 μl of the culture supernatant were injected and separated on the HPLC column using a previously published protocol (Geraci et al., 2022): Briefly, for separation of gluconate, a flow rate of 0,2 ml/min and a linear gradient of 25 min, reducing the concentration of buffer A from 82 % to 42 % was used. Before (5 min) and after (10 min) the gradient, the column was equilibrated with 82 % buffer A.

The mass-to-charge (m/z) ratios of the separated samples were analysed by MS operated in negative ion-mode with a mass range of 85-900 m/z. An extracted ion chromatogram of gluconate based on the exact mass of (M-H) ^−^ = 195,05 m/z (± 0,02) was generated and used to calculate the areas under the curve (AUC). The estimation of the amounts of gluconate contained in the samples was ensured by an appropriate calibration.

## Acknowledgement

The author’s work was supported in former times by the Deutsche Forschungsgemeinschaft. I thank Karl Forchhammer for generous hospitality, Heike Brötz-Österhelt for strains and Marina Borisova and Christoph Mayer for strains and many stimulating discussions. Special thanks go to Vikram Alba (Max Planck Institute for Biology/Protein Evolution, Tübingen) for modelling the PQQ-PqqU interaction. We thank Libera Lo Presti (Tübingen) for very helpful suggestions.

## Supplement

**Fig.S1:**
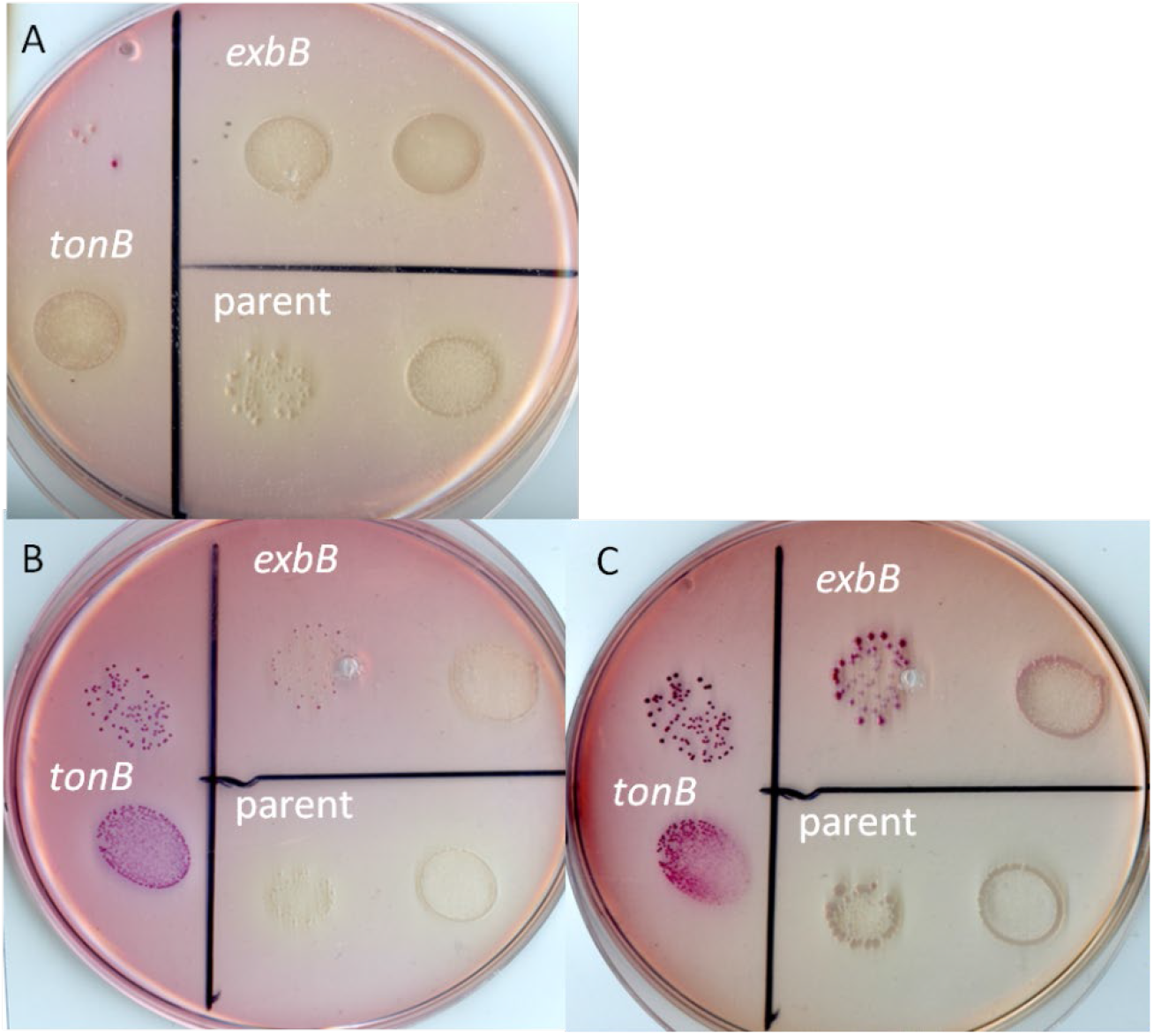
MaConkey agar without sugar. (A) supplemented with 10 μmol/l Fe(III)Na-EDTA; (B) without supplement; (C) the same plate 3 days later. Acid secretion due to iron limitation is demonstrated by the red color of the strains with impaired siderophore uptake systems.

