## Supplementary material for "The TonB dependent uptake of pyrroloquinoline-quinone (PQQ) and secretion of gluconate by *Escherichia coli* K-12": Fig.S1: Supplement.docx

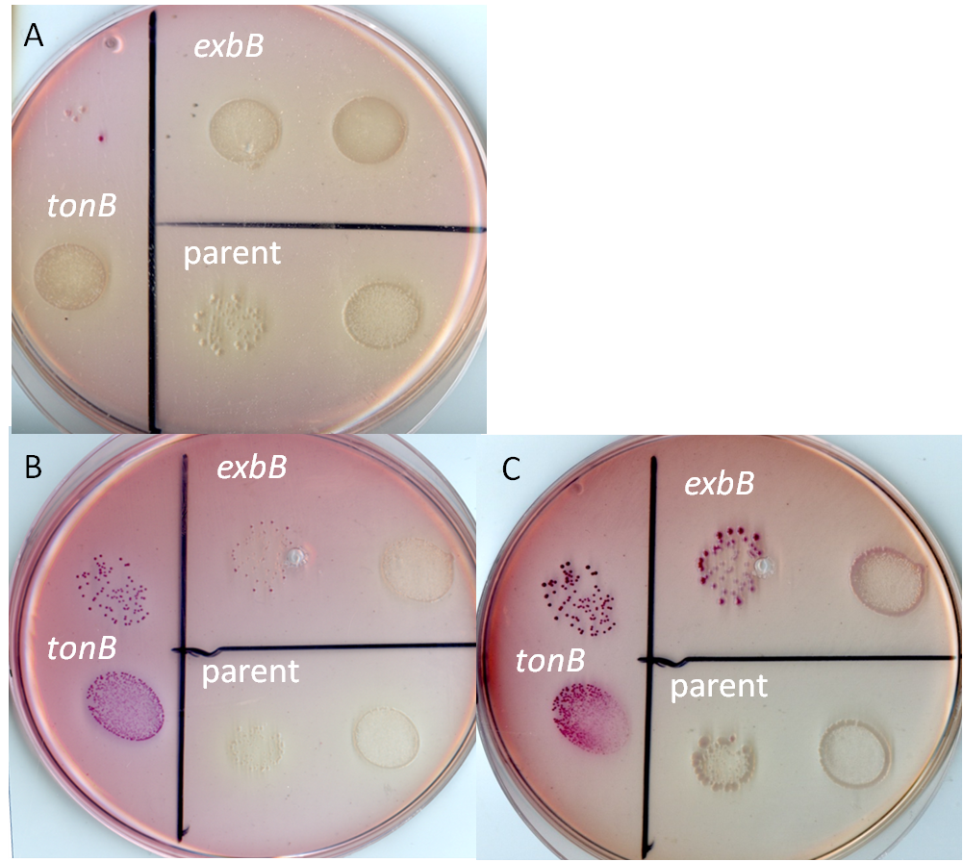

**Fig.S1: MaConkey agar without sugar**

(A) supplemented with 10 µmol/l Fe(III)Na-EDTA; (B) without supplement; (C) the same plate 3 days later. Acid secretion due to iron limitation is demonstrated by the red color of the strains with impaired siderophore uptake systems.
